# Array-based sequencing of filaggrin gene for comprehensive detection of disease-associated variants

**DOI:** 10.1101/103416

## Abstract

The filaggrin gene (*FLG*) is essential for skin differentiation and epidermal barrier formation with links to skin diseases, however it has a highly repetitive nucleotide sequence containing very limited stretches of unique nucleotides for precise mapping to reference genomes. Sequencing strategies using polymerase chain reaction (PCR) and conventional Sanger sequencing have been successful for complete *FLG* coding DNA sequence amplification to identify pathogenic mutations but this time-consuming, labour intensive method has restricted utility. Next-generation sequencing (NGS) offers obvious benefits to accelerate *FLG* analysis but standard re-sequencing techniques such as oligoprobe-based exome or customized targeted-capture can be expensive, especially for a single target gene of interest. We therefore designed a protocol to improve *FLG* sequencing throughput using a set of *FLG*-specific PCR primer assays compatible with microfluidic amplification, multiplexing and current NGS protocols. Using DNA reference samples with known *FLG* genotypes for benchmarking, this protocol is shown to be concordant for variant detection across different sequencing methodologies. We applied this methodology to analyze cohorts from ethnicities previously not studied for *FLG* variants and demonstrate usefulness for discovery projects. This comprehensive coverage sequencing protocol is labour-efficient and offers an affordable solution to scale up *FLG* sequencing for larger cohorts. Robust and rapid *FLG* sequencing can improve patient stratification for research projects and provide a framework for gene specific diagnosis in the future.

## INTRODUCTION

The filaggrin gene (*FLG*) encodes profilaggrin, a major epidermal structural protein with multifunctional roles essential for effective skin barrier formation (Sandilands et al. 2009; McAleer and Irvine 2013). Loss-of-function (LoF) variants in *FLG* have been identified as the causative genetic defect in the dry skin condition ichthyosis vulgaris (IV; OMIM 146700) and represent the most significant genetic variants associated with risk of atopic dermatitis (AD) (Palmer et al. 2006; Smith et al. 2006; van den Oord and Sheikh 2009). IV is one of the most common inherited skin diseases with surveys reporting a population prevalence of up to 1 in 250, with AD often present as a secondary trait but with lower penetrance (Wells and Kerr 1966; Smith et al. 2006). AD is a prevalent inflammatory skin barrier deficiency affecting up to 25% of children in developed countries (Tay et al. 2002; Sandilands et al. 2006; Odhiambo et al. 2009). The incidence of *FLG* LoF alleles in AD patients varies considerably around the world with Asian ethnic populations suggesting a 20 – 30% mutation burden whilst in northern Europe the incidence can be up to 50% (Sandilands et al. 2007; Brown et al. 2008a; Chen et al. 2011; Kono et al. 2014). The reasons underlying geographical variations in combined mutation frequencies are unclear but their maintenance within the gene pool is consistent with a heterozygote advantage (Irvine and McLean 2006; Thyssen et al. 2014; Eaaswarkhanth et al. 2016), although the basis for this is still under debate. Recent research has confirmed that *FLG* LoF variants are associated with early-onset, more severe and persistent AD, highlighting a primary skin barrier dysfunction in this disorder (Brown et al. 2008b; Fallon et al. 2009).

Distinct sets of LoF variants are present in different ethnicities. In the northern European population, two predominant sequence variants make up 80% of the mutation burden (Sandilands et al. 2007). In contrast, in east Asian ethnicities (cohorts from Japan, Taiwan, Singaporean Chinese, Korea and China), more diversity of *FLG* LoF variants are found with fewer predominating variants, resulting in a wider mutation spectrum compared to Caucasian populations (Chen et al. 2011; Irvine et al. 2011). In addition, many rare family-specific and country-specific LoF variants have been reported, adding to the global complexity of *FLG* pathogenic mutation identification. Compilation of published data has identified 85 *FLG* LoF disease-associated variants to date (Supplemental Table 1) and 252 *FLG* LoF variants identified in multiple ethnic populations (without disease information) from exome sequencing data deposited in the Exome Aggregation Consortium (ExAC; Supplemental Table 2) (Thyssen et al. 2014; Lek et al. 2016).

**Table 1.** *FLG* LoF variants detected in 14 samples analyzed with Sanger-sequencing, Roche 454 GS FLX+ and Illumina MiSeq sequencing platforms. Concordant mutation detection was largely achieved with all 3 NGS strategies with only the MiSeq platforms able to detect all *FLG* LoF variants in the 14 samples analyzed. ND = Known variant was not discovered; “-“ = No pathogenic variant identified.

| ID | Sanger sequencing | Roche 454 GS FLX+ | Illumina MiSeq 2 x 300 assay | Illumina MiSeq 2 x 250 assay |
| --- | --- | --- | --- | --- |
| IA-P003 | p.S1515X | p.S1515X | p.S1515X | p.S1515X |
| IA-P009 | - | - | - | - |
| IA-P014 | - | - | - | - |
| IA-P017 | p.E2422X | p.E2422X | p.E2422X | p.E2422X |
| IA-P021 | p.S406X;<br>c.6950_6957del8 | p.S406X; ND* | p.S406X;<br>c.6950_6957del8 | p.S406X;<br>c.6950_6957del8 |
| IA-P024 | c.1640delG | c.1640delG | c.1640delG | c.1640delG |
| IA-P025 | p.Q368X;<br>c.3321delA | p.Q368X;<br>c.3321delA | p.Q368X;<br>c.3321delA | p.Q368X;<br>c.3321delA |
| IA-P028 | c.7945delA | c.7945delA | c.7945delA | c.7945delA |
| IA-P062 | p.Q2417X | p.Q2417X | p.Q2417X | p.Q2417X |
| IA-P063 | c.2952delC | c.2952delC | c.2952delC | c.2952delC |
| IA-P083 | c.9040_9058dup19 | p.Q1790X; ND* | c.9040_9058dup19;<br>p.Q1790X | c.9040_9058dup19;<br>p.Q1790X |
| IA-P084 | p.S1302X; ND | p.S1302X;<br>p.S1515X | p.S1302X;<br>p.S1515X | p.S1302X;<br>p.S1515X |
| IA-P090 | c.4004del2 | c.4004del2 | c.4004del2 | c.4004del2 |
| IA-P094 | c.2282del4;<br>p.R2447X | c.2282del4; ND* | c.2282del4;<br>p.R2447X | c.2282del4;<br>p.R2447X |

\* ND allele frequency below 5%

**Table 2.** Summary of patient demographics and clinical features of the 3 ethnicities studied for *FLG* LoF variants. SCORAD (SCORing AD) index and objective SCORAD (oSCORAD) are used for clinical phenotyping. Complete metadata is provided in Supplemental Table 7 and Supplemental Table 10.

|  | Chinese | Malay | Indian |
| --- | --- | --- | --- |
| Subjects demographics |  |  |  |
| Subjects (*n*) | 279 | 36 | 19 |
| Age, mean (range) in years | 18.5 (2-70) | 22.4 (7-45) | 23.6 (8-60) |
| Male:female (*n*) | 202:77 | 28:8 | 12:7 |
| Ichthyosis vulgaris |  |  |  |
| No ichthyosis vulgaris (*n*) | 103 | 12 | 3 |
| Mild (*n*) | 80 | 13 | 6 |
| Moderate-Severe (*n*) | 84 | 11 | 8 |
| Not recorded (*n*) | 12 | 0 | 2 |
| Number of IV patients with *FLG* LoF (%) | 64 (39.0) | 7 (29.2) | 8 (57.1) |
| Atopic dermatitis |  |  |  |
| Mean Total SCORAD (+/- SD) | 47.2 (± 17.3) | 50.6 (± 19.7) | 50.6 (± 13.9) |
| Mean oSCORAD | 37.6 (± 15.0) | 38.8 (± 17.4) | 39.4 (± 11.4) |
| Mild AD / oSCORAD <15 (*n*) | 14 | 3 | 1 |
| Moderate AD / oSCORAD 15-40 (*n*) | 151 | 16 | 8 |
| Severe AD / oSCORAD >40 (*n*) | 110 | 17 | 10 |
| IV only, AD not reported (*n*) | 4 | 0 | 0 |
| Mean oSCORAD of samples with *FLG* LoF | 37.5 (± 15.3) | 38.9 (± 17.6) | 38.2 (± 8.6) |
| Mean oSCORAD of samples with WT *FLG* | 37.6 (± 15.0) | 38.8 (± 17.4) | 40.0 (± 11.3) |
| AD Age of onset<5 years age (*n*) | 143 | 13 | 8 |
| AD Age of onset>5 years age (*n*) | 124 | 23 | 11 |
| AD Age of onset unknown (*n*) | 12 | 0 | 0 |

The filaggrin gene is a member of the S100 fused-type gene family, a group of genes with a conserved exon structure and likely shared functions in epithelial biology (Gan et al. 1990; Henry et al. 2012; Strasser et al. 2014). *FLG* transcription is initiated in exon 2 with the majority of the profilaggrin polyprotein produced from the extremely long (>12Kb) and highly repetitive exon 3. The genetic architecture of exon 3 consists of 10, 11 or 12 tandemly arranged filaggrin repeat units, each approximately 972 nucleotide base pairs (bp) long and joined by a 21 bp linker region (Gan et al. 1990). Intragenic copy number variation (CNV) of filaggrin repeats increases the gene structure complexity, with each repetitive unit almost identical in sequence (Sandilands et al. 2007; Sasaki et al. 2008; Brown et al. 2012). The presence of these numerous highly repetitive stretches of nucleotide sequences makes primer design challenging, and restricts primer site availability to a small number of unique bases present in the filaggrin repeat units and their linker regions (Sandilands et al. 2007; Sandilands et al. 2009). Taken together, the genetic architecture of *FLG* makes routine analysis by PCR and Sanger sequencing extremely laborious.

To study genetics of AD properly in any given population, appropriate stratification to determine risk predictions is essential, whether for research studies or clinical trials. Therefore a fast, robust, comprehensive and affordable method for *FLG* coding DNA sequence (CDS) analysis is required. To date, most *FLG* studies have taken the approach of targeted screening for the predominant common LoF variants to report allele frequencies after full sequencing of an initial small batch of test patient samples. This strategy runs the risk of under-reporting in populations where *FLG* LoF variantss are highly diverse, due to discovery sampling effects (Clark et al. 2005). Whilst full Sanger sequencing of the *FLG* CDS is not feasible in all laboratories, it is still necessary in order to provide comprehensive study results (Sandilands et al. 2007). Target capture techniques such as exome sequencing captures *FLG* LoF variants but is currently expensive if the aim is to study a single gene. Also, due to the occurrence of rare and *de novo* family specific *FLG* LoF variants, the over reliance on publically available aggregated summary sequencing data will not inform sufficiently for LoF targeted screening of *FLG* (Margolis et al. 2014; Lek et al. 2016; Ruderfer et al. 2016).

In this paper we describe a simple, robust and cost-effective PCR-based method for analyzing the entire CDS of *FLG* including the well known intragenic CNVs. This protocol uses microfluidics technology to reduce the amount of DNA starting material needed, increases sample throughput and reduces the required operator time for comprehensive *FLG* genotyping. Coupling this sample preparation protocol with relatively long NGS reads and a streamlined bioinformatics pipeline for analysis, we present a scalable solution for efficient cohort analysis. This protocol also removes the ascertainment bias introduced by only screening for the most common population-specific *FLG* LoF variants (Clark et al. 2005). The potential usefulness of a robust protocol to identify *FLG* sequence variants extends beyond the research environment and could aid clinical diagnosis and patient treatments as more pharmacogenetic therapies for AD are developed (Brown and McLean 2012; Margolis et al. 2013). This approach to sequencing *FLG* could also be applied to other large complex genes that are difficult to analyze, such as the additional members of the S-100 family.

## RESULTS

### Establishing a validated protocol for *FLG* CDS amplicon sequencing

We designed and optimized a set of 48 *FLG*-specific primer assays (containing sequencing adapters) to span the entire *FLG* CDS and generate overlapping amplicons suitable for downstream processing and paired-end sequencing with Illumina MiSeq NGS (MiSeq Reagent Kit v2, 2 × 250 bp read mode; Supplemental Table 3). Each of the 48 primer assays was designed with an amplicon size between 400 and 500 bp. Analyzing conventional PCR reactions confirmed accurate amplicon sizing of a single major PCR band with minimal generation of non-specific products (Figure 1A). Overlapping primer pair assays were designed to provide redundancy in sequencing reads across primer binding sites. In particular, the regions between repeat 7 and repeat 8 or 8^1^ were heavily overlapped with amplicon design to improve coverage (Figure 1B). This also enabled confirmation that single nucleotide variations (SNV) were not present in primer binding sites, as this could result in the loss of PCR amplicons.

**Table 3.** 18 *FLG* LoF variants identified in Malay and Indian IV and/or AD cohorts using MiSeq 2 × 250 bp protocol. Singaporean Malay and Indian ethnicities have both unique and recurrent *FLG* LoF variants compared to other published studies (see Supplemental Table 1). * = not previously reported LoF variant in AD related published literature.

| ID | *FLG* mutation | Ethnicity | dbSNP ID | South Asian ExAC MAF | East Asian ExAC MAF |
| --- | --- | --- | --- | --- | --- |
| 1 | p.R501X | Indian | rs61816761 | 21/16512 | 0/8654 |
| 2 | p.S507X * | Malay | - | Not reported | Not reported |
| 3 | c.2282del4 | Malay | rs558269137 | 122/16510 | 0/8654 |
| 4 | c.3321delA | Malay | rs200519781 | 0/16512 | 82/8654 |
| 5 | p.R1140X | Indian | - | 16/16512 | 0/8652 |
| 6 | c.4812ins5 * | Indian | - | Not reported | Not reported |
| 7 | c.5024delC * | Indian | rs749542190 | Not reported | Not reported |
| 8 | c.5187delA * | Indian | - | Not reported | Not reported |
| 9 | c.5192_5199dup8 | Malay | rs754949514 | Not reported | Not reported |
| 10 | p.Q2123X * | Indian | rs145119684 | 8/16512 | 0/8654 |
| 11 | c.6834del5 | Malay | rs772007167 | Not reported | Not reported |
| 12 | c.6950del8 | Malay; Indian | rs578184315 | Not reported | Not reported |
| 13 | p.S2344X * | Malay, Indian | rs372754256 | Not reported | Not reported |
| 14 | c.7333delC | Indian | - | Not reported | Not reported |
| 15 | p.R2447X | Malay, Indian | rs138726443 | 35/16512 | 1/8652 |
| 16 | c.7487delC * | Indian | rs375277670 | 0/16512 | 5/8650 |
| 17 | p.R2613X * | Indian | rs567795279 | Not reported | Not reported |
| 18 | c.8088delG | Malay | - | Not reported | Not reported |

**Figure 1.**
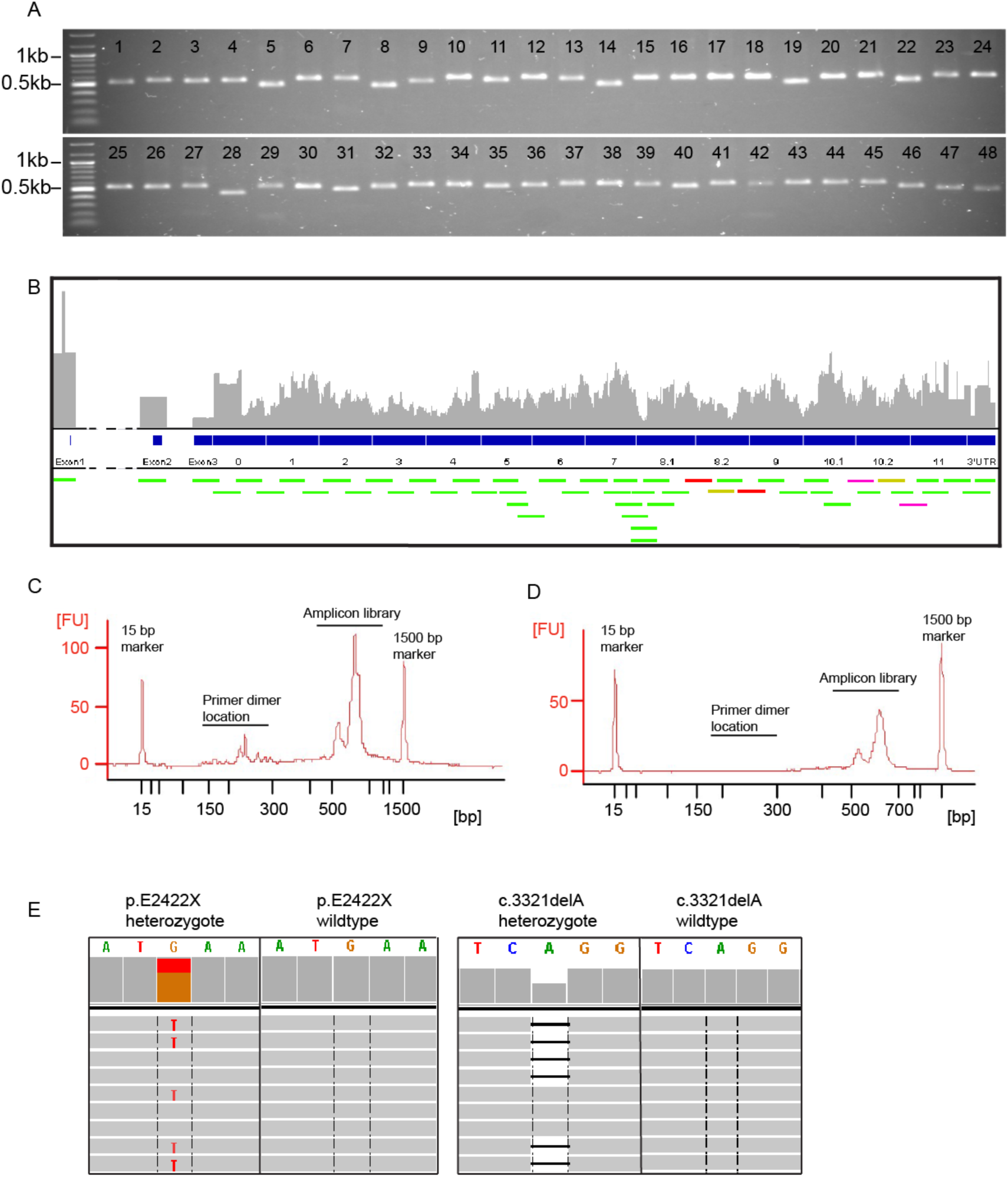
Primer validation, nucleotide base coverage and library pooling for *FLG* CDS sequencing. (A) 48 primer assays for the MiSeq 2 × 250 bp protocol were validated by PCR and gel electrophoresis to confirm expected amplicon size and absence of non-specific products. (B) Amplicons were designed with overlapping coverage and are visualized across our in-house 12-repeat *FLG* CDS (green bars). Primer assay 34 (yellow bars), 35 (red bars) and 41 (purple bars) produce multiple amplicons when there are intragenic duplication of repeats 8 and 10. (C) Electropherogram of a representative pre-cleanup sample library on the Agilent Bioanalyzer showing the presence of primer dimers. Single sample amplicon library with a peak between 550 and 650 bp after the addition of adaptor tags for sequencing. (D) Electropherogram of the post-clean up final library pool containing amplicons from 92 samples without primer dimers. (E) Integrated Genome Viewer screenshot showing a heterozygous *FLG* nonsense LoF variant carrier (p.E2422X) compared to wildtype and (F) a heterozygous *FLG* frameshift LoF variant carrier (c.3321delA) compared to wildtype.

The Access Array 48.48 integrated fluidic circuit (IFC) chip (Fluidigm, US), hereafter referred to as the Access Array IFC, allows for parallel amplification of 48 amplicons for 48 different DNA samples, simultaneously generating 2304 PCR reactions with nanolitre volumes suitable for downstream massively parallel sequencing projects (Lange et al. 2014). 96 different DNA samples were analyzed in a pilot experiment using two Access Array IFC chips. Amplicons were tagged with sample-specific barcodes during the PCR process to facilitate downstream multiplexed MiSeq NGS (Illumina). Amplicon pools from each Access Array IFC were visualized for correct amplification product size with Bioanalyser (Agilent) prior to bead purification (Figure 1C). Out of the 96 samples run on the two Access Array IFC, four samples did not produce libraries and were omitted from further analysis. Subsequently the PCR products from two Access Array IFC chips were combined to produce a single amplicon pool containing the 92 barcoded DNA amplicon libraries that were successfully amplified. These 92 were again sized with Bioanalyser (Figure 1D), before sequencing on the Illumina MiSeq using 2 × 250 bp mode (see Methods). Importantly, primer dimer contaminants that could potentially interfere with downstream sequencing were removed from the final amplicon pool using bead purification (Figure 1D). Sequencing of this pool on the Illumina MiSeq 2 × 250 bp mode resulted in high average sequencing depth per base that corresponds to the number of multiplexed DNA samples.

Sequence reads were mapped to the previously reported 12-repeat *FLG* reference sequence (Sandilands et al. 2007) using the BWA-MEM algorithm (Li 2013). This reference sequence contains the variably present additional intragenic CNV allelic variants of repeat 8 and repeat 10 but is not currently documented in the NCBI RefSeq database. The mapped reads were processed with the GATK suite of tools (McKenna et al. 2010). Quality control of the reads and variants was ensured by the internal functions of GATK (see Methods). By default our pipeline returns mapping files (BAM format) and variant call files (VCF format). Functional annotation of variants was added with the SnpEff tool (Cingolani et al. 2012). Visual inspection of identified null-mutations was performed using the Integrated Genome Viewer (Broad Institute; Figure 1E) (Robinson et al. 2011). In an optional pipeline step, the coordinates of identified variants can also be converted to the *FLG* gene 10-repeat reference sequence that is found in current genome annotations (RefSeq GRCh37 or GRCh38).

One of the genomic features of *FLG* that we aimed to assess in our samples was the presence or absence of intragenic CNV of repeats 8 and 10 (Figure 2A), as this is an important risk factor for AD (Brown et al. 2012). Complete 12-repeat *FLG* CDS coverage was achieved with 48 primer assays that encompassed the known intragenic CNV allelic variants. Primer assays 34, 35 and 41 (Figure 1B; Supplemental Table 3) amplify multiple segments of *FLG* repeats 8 and 10 and their CNVs. Amplicons were then mapped to the CNV repeat regions using unique nucleotide bases that differentiate between the *FLG* repeats.

**Figure 2.**
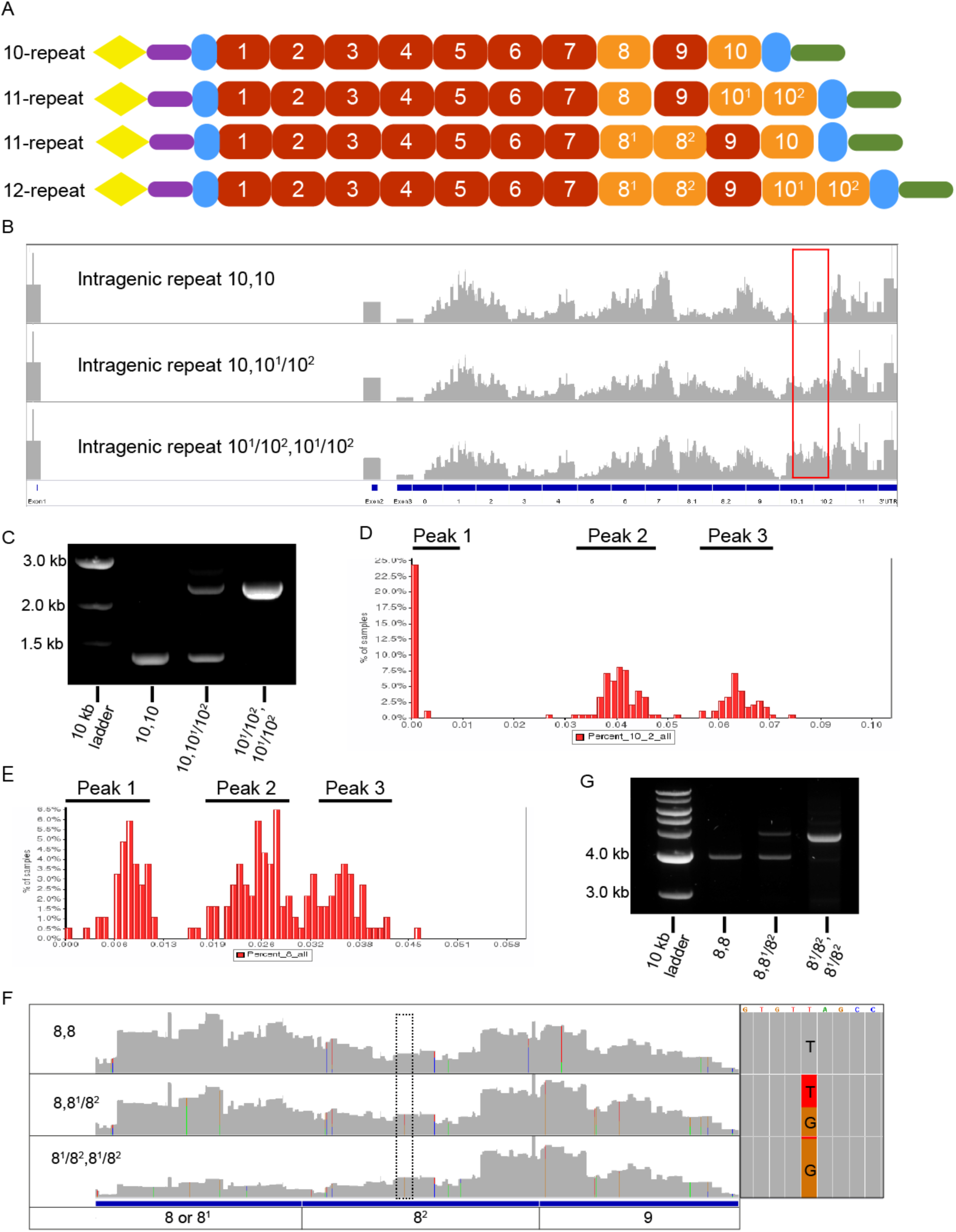
Identification of copy number variation (CNV) in the *FLG* gene structure. (A) Diagram showing *FLG* alleles generated with combinations of intragenic CNVs. Yellow triangle = S100 domain; purple bar = B-domain; red square = *FLG* repeats; orange square = intragenic CNV *FLG* repeats; blue rectangle = partial *FLG* repeats; green bar = C-terminal domain. (B) IGV plot of sequencing reads showing the presence or absence of coverage in the repeat 10 region (red box). Read coverage is used for estimating allele size with regard to CNV in overall gene structure. Repeat 10 CNV allelic genotype is indicated on each individual IGV plot. (C) Estimations for repeat 10 CNV were confirmed with corresponding conventional PCR and gel electrophoresis. Repeat 10 CNV allelic genotype is indicated below each lane. (D) Coverage predictions were generated with read coverage ratios to generate distribution graph for multiple samples (*n*=92). Three clear peak clusters observed in the distribution graphs correspond to the intragenic repeat 10 CNV predictions. Peak 1 corresponds to homozygous for repeat unit 10 only, peak 2 corresponds to heterozygous repeat 10 units - allele 1 contains 10; allele 2 contains 10^1^/10^2^. Peak 3 on the distribution graph corresponds to homozygous CNV repeat 10 units 10^1^/10^2^. (E) For repeat 8 CNV, coverage predictions were generated using reads coverage ratios. Distribution plot indicating 3 peak clusters corresponding to homozygous repeat 8 unit (Peak 1), peak 2 corresponds to heterozygous repeat 8 and 8^1^/8^2^ units and peak 3 on the distribution graph corresponds to homozygous repeat 8^1^/8^2^ units on both alleles. (F) Repeat 8 CNVs were also determined by the presence of intra-allelic nucleotide variant in repeat 8^1^ (c.8673) and its corresponding position in repeat 8^2^ (c.9645 as indicated by the dotted box). (G) Repeat 8 CNV alleles were confirmed with conventional PCR and gel electrophoresis.

For identification of repeat 10 CNV, we observed that absence of duplication (i.e. having the canonical repeats 9, 10 and 11 in sequence) would coincide with a complete drop in coverage in the region 22500-23000 of the 12 repeat reference CDS, corresponding to the position of 10^2^ (Figure 2B). Likewise, the presence of repeat 10 duplication (i.e. repeats 9, 10^1^, 10^2^, 11) would manifest as the presence of sequencing reads covering this same region of the *FLG* reference CDS (Figure 2B). DNA sample profiles for repeat 10 CNV were confirmed using PCR and gel electrophoresis (Figure 2C). We were therefore able to calculate four ratios from our NGS data based on the number of mapped reads in the 10^2^ region normalising to: 1) total number of mappings, 2) total number of reads with at least one mapping, 3) number of reads in *FLG* coordinates for 10^1^ (15480-15950) and 4) 10^2^ (23500-24000), of the *FLG* 12-repeat reference sequence where coverage is stable over approximately 500 bp. The distribution of these ratios shows three distinct peaks: homozygous reference (near zero on the X axis), heterozygous and homozygous duplicated (Figure 2D). The latter two peaks can sometimes merge at the tails depending on the quality of the samples. If this is observed, samples in this overlapping region are marked as ambiguous. When three of these ratios are in agreement the repeat is called automatically (90 out of 92 samples). If not, visual inspection of the data with IGV is used to assess the repeat status (2 out of 92). These peaks can be slightly shifted for separate batches of samples but are always distinctly observable in the batches processed thus far.

Filaggrin repeat 8^1^ and 8^2^ have near identical sequence homology resulting in NGS reads potentially mapping to either genomic region. When calculating intragenic variation for filaggrin repeat 8 we studied coverage using the 4 ratios described above but did not rely on coverage alone to distinguish CNV identity (Figure 2E). In Singapore Chinese DNA samples that we analysed we could differentiate between repeat 8^1^ and 8^2^ using a T to G intragenic allelic variant between CNV repeats (Figure 2F). However, this particular variant was not informative for CNV identification in Caucasian samples we analysed (data not shown). Based on the allele frequency of the SNV together with coverage data we make a call regarding the presence of repeat 8^2^. Finally, this result was correlated with a specific conventional PCR and electrophoresis to confirm the repeat 8 CNV profile in representative samples (Figure 2G).

The methodology described here constitutes a validated high throughput PCR-based amplification, NGS and bioinformatics analysis protocol for comprehensive analysis of the *FLG* CDS (Supplemental Figure 1). This protocol is amenable to further scaling up of sample throughput, thus providing a means for rapid turnaround of large cohorts for targeted *FLG* resequencing. Analysis of sequencing paired-end reads provided high quality *FLG* genotyping that can provide more than adequate depth for base calling in mutation analysis and also estimates of intragenic CNV.

### *FLG* LoF variant detection across multiple sequencing methodologies

Using the 92 DNA samples that were fully analyzed for the pilot experiments, we compared two additional *FLG* sequencing strategies with the 2 × 250 bp protocol, described above, for coverage, sequencing depth and mutation detection. These were (1) Roche 454 GS FLX+ pyrosequencing using primers with average length of 538 bp (range 267 – 758 bp) based on the Sandilands et al. (2007) protocol (Supplemental Table 4) and (2) an Illumina MiSeq reagent kit v3, 2 × 300 bp mode with overlapping amplicons designed with an average of 546 bp (Supplemental Table 5). Comparing the 3 sequencing strategies, an additional 4 samples failed on at least one platform and were therefore removed from further analysis. The remaining 88 DNA samples were used for subsequent cross-platform comparison. 14 of the 88 DNA samples had been previously fully Sanger sequenced for *FLG* CDS using well established published primers (Sandilands et al. 2007; Chen et al. 2011) and thus their *FLG* LoF variant profile was known. *FLG* variant profile in these 14 samples across the 3 investigated protocols was similar but only the MiSeq methods detected the full complement of disease associated *FLG* LoF variants (Table 1). In the additional 74 samples that were not Sanger sequenced, MiSeq methods detected 3 additional *FLG* LoF variants compared to Roche 454 GS FLX+ protocol even though coverage was sufficient (c.6950_6957del8, p.R2447X, c.9040_9058dup19; Supplemental Table 6). Regarding target base coverage, successfully run homozygous 12 repeat samples were covered by at least 100-fold read depth across 94.9% of exon bases for Roche 454 GS FLX+ assay, 99.3% on Illumina MiSeq 2 × 300 bp assay and 100% for Illumina MiSeq 2 × 250 bp assay.

The Roche 454 GS FLX+ protocol produced longer read lengths that potentially provided more accurate read mapping and better resolution of intragenic CNV repeats. However, although this method was able to identify the majority of *FLG* LoF variants correctly (Table 1), major difficulties were encountered in distinguishing true insertions and deletions from false positives due to the inherent limitation of the 454 GS FLX+ in interpreting homopolymer sequences (Balzer et al. 2011). Additionally, sequencing depth of *FLG* CDS lacked uniformity, probably due to uneven amplicon size within our design.

We used the MiSeq reagent kit v3, 2 × 300 bp protocol to maximize amplicon length with Illumina sequencing that potentiated more accurate mapping for this system and larger overlap of amplicons. However, the MiSeq reagent kit v3, 2 × 300 bp mode resulted in suboptimal quality control, reporting poor base call Q scores that raised the probability of false-positive variant calls in sequencing reads (63.6% of bases had a Q score of 30 or higher for the pilot run of 88 samples). Despite sequencing data with suboptimal Q30 scores, the erroneous variants were interspersed at low frequency throughout the gene, so that with deep sequencing per base coverage we were able to unambiguously identify all *FLG* LoF variants in line with the other sequencing techniques analyzed (Table 1). In comparison, primer assays specifically designed for Illumina MiSeq 2 × 250 bp protocol produced good overall sequencing coverage with higher quality scores (80.9% of bases with Q score >30 for pilot experiment of 92 samples), *FLG* LoF variants were all detected according to the known sample profiles (Table 1) and multiplexing 88 samples in one MiSeq lane gave >100 reads per base comprehensively covering 100% of the *FLG* CDS.

Protocols using Illumina MiSeq 2 × 300 and 2 × 250 bp modes were able to identify all *FLG* LoF variants identified from either Sanger or Roche 454 GS FLX+ sequencing but also identified additional clinically relevant LoF variants. Therefore the MiSeq protocol was used for subsequent studies and specifically the MiSeq reagent kit v2, 2 × 250 bp mode because of the base call quality.

### Validated *FLG* sequencing protocol applied to discovery cohorts highlight novel population specific disease associated *FLG* LoF variants

We used the Access Array IFC and Illumina MiSeq 2 × 250 bp sequencing protocol to analyze 279 Singapore Chinese IV and/or AD patient samples to obtain a comprehensive estimate of disease-associated LoF allele frequency from fully sequenced *FLG* (Table 2; Figure 3; Supplemental Table 7). 85 of these IV and/or AD samples had previously been Sanger sequenced for the entire *FLG* CDS in our earlier study (Chen et al. 2011). This reanalysis provided a validation experiment for *FLG* LoF variant detection in comparison to Sanger sequencing. In this comparison we identified one additional mutation with Access Array IFC amplification and MiSeq sequencing that was not picked up in our previous study (Supplemental Table 8). Complete CDS sequencing of the remaining 194 DNA samples identified 11 additional variants in the Singapore Chinese population, bringing the total number of variants identified to 32 (an increase from 22 in our previous study), with five variants not previously reported in the literature (Figure 3; Figure 4). Fisher’s exact test on the *FLG* LoF variants using ExAC (version 0.3.1) exome data as population controls showed that 14 LoF variants reached individual significance of p<0.05 (Supplemental Table 9).

**Figure 3.**
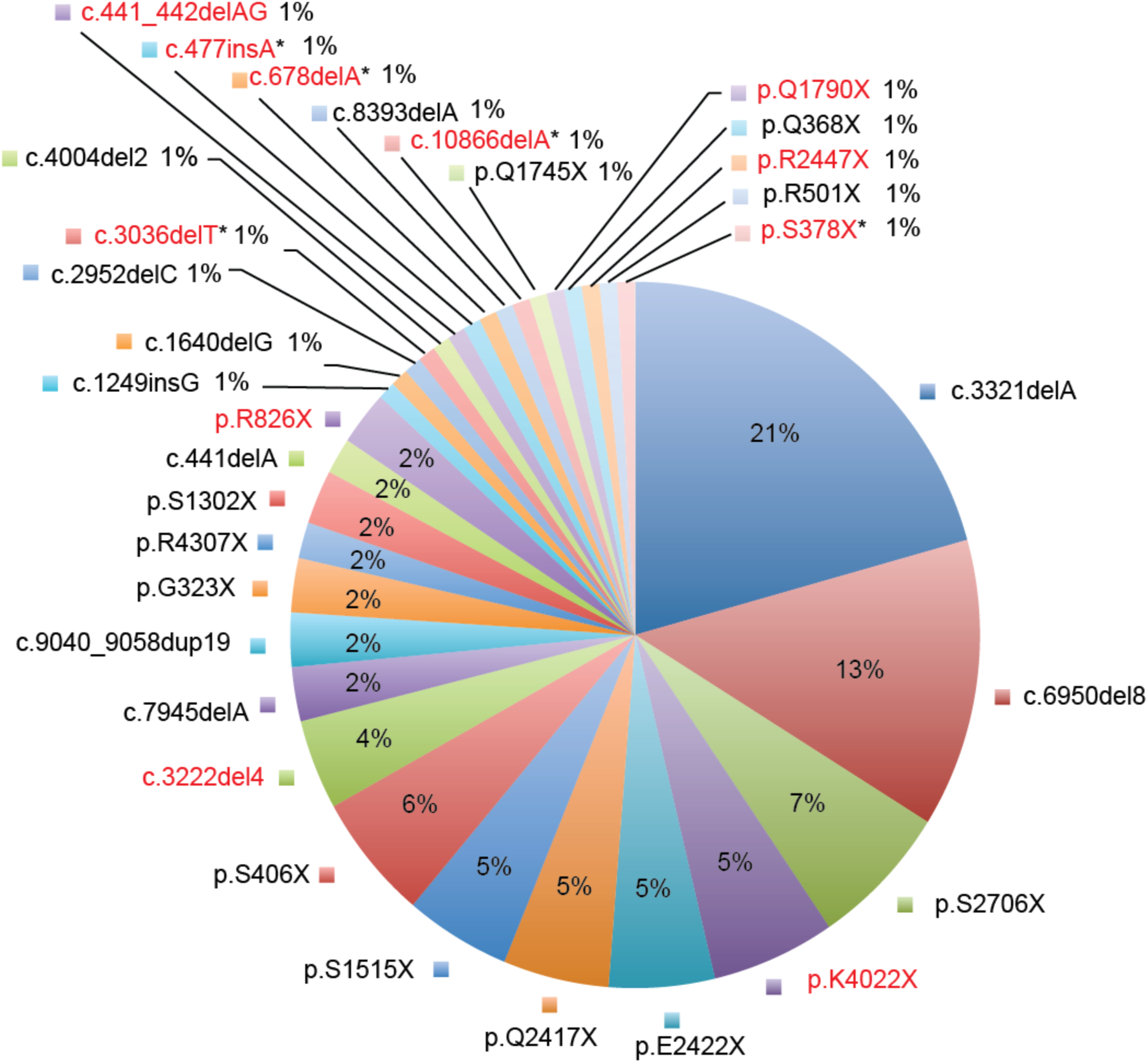
Spectrum of disease associated *FLG* LoF variants identified in the Singaporean Chinese IV and/or AD population with MiSeq 2x250bp workflow. *FLG* CDS of 279 Chinese patients were fully sequenced to identify *FLG* LoF variants. Chart shows the percentage contribution of 32 pathogenic variants identified in the Singaporean Chinese cohort. 11 additional variants (red text) were identified in this population compared to our previous survey (Chen et al. 2011), of which 5 variants have not been previously reported in the literature (marked with *).

**Figure 4.**
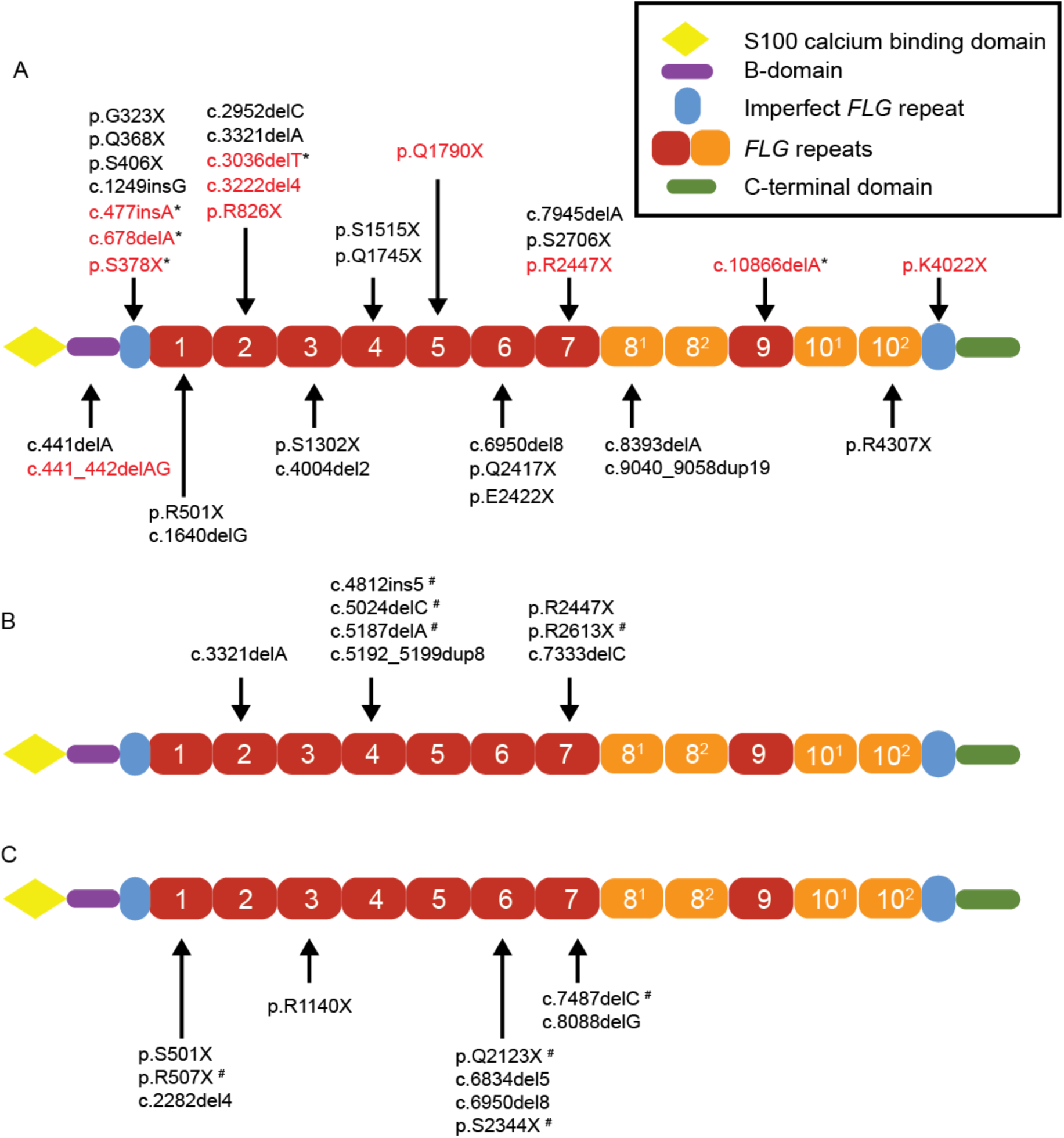
Schematic of the profilaggrin molecule showing the domain position of LoF *FLG* variants from Singaporean patients with IV and/or AD. (A) Positions of *FLG* LoF variants from Singapore Chinese samples. *FLG* variants not previously reported in Singapore Chinese samples are highlighted in red. Novel LoF variants not previously reported are marked with * (B) Position of *FLG* LoF variants identified in this study from Indian and (C) Malay samples from Singapore. LoF variants not previously identified in AD patient-based studies marked with ^#^.

Next, we analysed IV and/or AD patients in Singapore from Indian and Malay ancestry, two Asian ethnic populations that have not been previously analysed for disease-associated *FLG* variants. We sequenced 19 Singapore Indian patients and 36 Singapore Malay patients using Access Array IFC and MiSeq sequencing protocol (Table 2; Supplemental Table 10). We identified *FLG* LoF variants in nine of the Indian samples (Table 3) and nine Malay samples (Table 3), therefore predicted to have disease relevance, as reported for all *FLG* LoF variants in other ethnic groups. All predicted disease-associated sequence variants were analysed for predicted function using MutationTaster2 (Schwarz et al. 2014) and validated by Sanger sequencing (Sandilands et al. 2007).

Applying the Access Array IFC *FLG* sequencing methodology to cohorts of divergent ethnic ancestry, we have analyzed a total of 334 Singaporean IV and/or AD samples. Specifically using MiSeq 2 × 250 bp protocol we identified additional *FLG* LoF variants, bringing the combined mutation allele frequency for Singapore Chinese AD patients up to 32.3% (analysis of 279 AD patients). *FLG* LoF variants were also investigated in Indian and Malay ethnicities from Singapore with IV and/or AD, which identified additional LoF variants and confirms a variable incidence of *FLG* linked AD between different ethnic groups (Indians at 47.4% in 19 patients and Malays 25% in 36 patients; Figure 4). The presence of a *FLG* LoF variant in patients did not significantly alter the SCORAD from those without *FLG* LoF variants in the cohorts studied (Table 2).

## DISCUSSION

The association of *FLG* LoF variants with skin and allergic diseases has been one of the most profound and impactful discoveries in human molecular genetics. Mutations in *FLG* have an impact both on causation and on risk of disease that blur the boundaries between single gene and complex traits disorders (Palmer et al. 2006; Smith et al. 2006; Irvine et al. 2011). Various hypotheses have been put forward about the potential evolutionary advantage of maintaining *FLG* LoF alleles in populations, ranging from vitamin D synthesis to trans-dermal immunisation, although no clear advantage has been confirmed (Irvine and McLean 2006; Thyssen et al. 2012; Thyssen et al. 2014; Nagao and Segre 2015; Eaaswarkhanth et al. 2016). Due to the size and complexity of the *FLG* gene, together with the cost and analysis time, the complete *FLG* CDS is rarely comprehensively sequenced and therefore the full spectrum of *FLG* variants is not completely realized in many patient cohorts. A non-NGS ‘*FLG*-shotgun’ sequencing method has previously been reported but not widely adopted possibly due to bacterial cloning steps (Sasaki et al. 2008). The capacity to efficiently study *FLG* is important because of its role in the pathogenesis of common skin disorders such as IV (1 in 250) and AD (1 in 5) that could be targeted with early therapies, such as emollients, to prevent disease progression. In this study we describe a protocol for amplification and sequencing of the entire *FLG* CDS for multiple patients with a semiautomated and affordable methodology that costs approximately 10 times less than exome target capture. We also show that this protocol is robust, reliable and has potential for scalable sample preparation for small and large research projects or clinical studies.

NGS has changed the landscape of mutation discovery with whole exome sequencing (Ng et al. 2009) and target capture of candidate genes or genetically linked regions becoming a common laboratory practice for research and diagnostic outcomes (Berg et al. 2011; Goudie et al. 2011; Artuso et al. 2012; Scott et al. 2012). In the work presented here, we used the power of NGS but circumvented the challenges of only resequencing one target gene by incorporating a multiplexed PCR step that provides an affordable way to look at *FLG* CDS in multiple samples in parallel. The relatively small capture size of 15 Kb for targeted re-sequencing experiments provided the opportunity to design an overlapping PCR strategy for *FLG* with inbuilt redundancy to capture the entire CDS. This, coupled with use of the Access Array IFC, facilitates high-throughput PCR amplification of multiplexed samples. This approach requires minimal amounts of high quality starting DNA for the assay (50 ng), plus reduced PCR reagents per reaction using the microfluidics set up (0.16 μl), and significantly reduced manpower is required in comparison to other strategies, making this an extremely cost-efficient protocol to run. Flexibility in the number of pooled DNA sample amplicon libraries prepared for a single MiSeq sequencing lane is also an advantage and can be tailored to experimental size and sequencing depth required - the upper limit currently determined by the availability of unique sample barcodes. We have routinely multiplexed 96 samples, but have also successfully multiplexed and analyzed 192 DNA samples on one Illumina MiSeq lane. Extrapolating from our current results, 384 samples could be successfully sequenced in one MiSeq lane with sufficient depth and full CDS coverage for mutation detection.

The benefits of our assay over conventional PCR strategies are clear, but there are potential problems and pitfalls to be considered. Sample batching can result in time delays prior to running the assay, although it is possible to run less than 96 samples albeit at increasing cost per sample. Experimentally there is always the possibility of amplification problems using primer assays - for example, the presence of an SNV in a primer-binding site of one allele could result in primer-binding failure, or reduced efficiency in generation of that allele’s amplicon during PCR amplification (allele dropout). Detection of allele dropout remains challenging and although we have not detected this in our sequencing so far, PCR amplification methods can be subject to loss of coverage of one allele. To reduce the chances of this happening, we designed all our primer assays with overlapping amplicon fragments and can check for SNVs in the primer sites if low sequencing read coverage is detected. Other potential pitfalls are mapping errors on this highly repetitive gene and reference bias. We retained standard calling thresholds for GATK for detection of mutations that allow for considerable deviation from the theoretical allele frequencies of 0%, 50% and 100%. This potentially leads to an increased number of false-positives, but conversely we were able to detect a heterozygous variant (c.9040_9058dup19) at 11% mutant to 89% wild type ratio. We therefore validated all deleterious variants with conventional PCR and Sanger sequencing as an additional confirmation step to ensure accurate reporting.

There are also practical cost considerations in the initial set-up. The initial outlay for this assay requires high quality primer assays and specialist equipment for NGS, and access to a Fluidigm thermocycler is essential for running the Access Array 48.48 IFC. We have designed this assay to run on the Illumina MiSeq for entry-level costing and it is closely related to the MiSeqDX that is increasingly found in clinical units and has been approved by the FDA for clinical diagnostics for certain applications. The MiSeq outputs fewer reads than other Illumina systems but offers more flexibility regarding batch size (48 to 384 samples depending on price point and sample throughput). Operator time needed to complete large batches of *FLG* sequencing is significantly reduced compared to Sanger sequencing. 96 samples can be set up on the Access Array IFC and then run on the Fluidigm thermocycler in one day; subsequently sample QC and pooling before MiSeq takes an additional day followed by MiSeq sequencing run according to local turnover time. Finally, mapping and variant calling can be completed in a few hours depending on the available computational capacity. Optional Sanger sequencing validation requires further time determined by local research environment constraints. As a working example, complete *FLG* CDS screening for an entire batch of 96 samples can be completed with a conservative time estimate of 4 weeks in our research environment.

In conclusion, we present a fast and robust method to approach high-throughput sequencing of the long and highly repetitive *FLG* coding sequence. This validated protocol enables rapid identification of gene mutations in patients to facilitate research studies and possibly genetic diagnostics in the future. Illumina sequencing platforms are widely used for clinical diagnostics, and therefore we envisage this as a step toward developing a clinical diagnostic strategy for *FLG* sequencing. Making *FLG* genotyping accessible to researchers and clinicians studying patients from any ethnicity with an unbiased approach is vital to progress a precision medicine approach to AD and indeed any other difficult to sequence genes. The identification of *FLG* risk genotypes will allow for a more informed approach towards treating and hopefully preventing the development of AD. The heterogeneous nature of AD and the genetic basis of inter-individual differences in drug responses will make risk genotypes particularly relevant for the development of personalised pharmacogenetic therapies and the use of skin barrier maintenance treatments.

## METHODS

### Sample collection

A total of 367 samples were used in this study (Supplemental Table 11). Within this, 334 patients were from Singaporean ethnicities diagnosed with IV and/or AD, recruited from National Skin Centre (NSC) Singapore by the specialist Eczema clinician team. All samples were collected with DSRB approval and in accordance with the Declaration of Helsinki. Clinical scoring was completed by trained clinicians using the well established SCORing AD (SCORAD) index developed by the European Task Force on Atopic Dermatitis (Oranje et al. 2007). Blood or saliva samples were collected with informed consent for DNA analysis; blood was processed fresh and saliva was stored at −30°C until processing. Genomic DNA was extracted from saliva samples using the Oragene DNA OG-500 kit (DNA Genotek Inc.) according to the manufacturer’s instructions. Genomic DNA from whole blood was extracted using the Nucleon Illustra™ BACC2 Genomic DNA Extraction Kit (GE Healthcare) according to the manufacturer’s instructions.

### Access Array 48.48 IFC target-specific primer design and validation

*FLG* primers were designed specifically for the different sequencing platforms tested. Primer assays with amplicons in the range of 400 – 500 bp were designed for Illumina Reagent Kit V2 MiSeq 2 × 250 bp sequencing. Target-specific primers for the Roche 454 GS FLX+ were designed based on the *FLG* primers used by the McLean group and gave overlapping PCR products with an average length of 538 bp (Sandilands et al. 2007; Chen et al. 2011). Partial redesign of *FLG* CDS PCR primers from (Sandilands et al. 2007) were used to produce amplicons with an average of 546 bp for use in Illumina MiSeq 2 × 300 bp sequencing.

*FLG* specific primers were validated according to the Access Array System User Guide for Illumina and Roche sequencing systems, these were termed the “inner” primers according to the user guide nomenclature (Fluidigm). All primers were designed to have universal adaptor sequences (CS1/CS2) attached to their 3’ ends. These sequences contained complementary sequences for amplicon barcoding. All forward primers contained universal adapter sequence ACACTGACGACATGGTTCTACA and all reverse primers universal adapter sequence TACGGTAGCAGAGACTTGGTCT.

Primers were validated using PCR workflow according to the Access Array manual (Fluidigm). PCR reactions (10μL) were conducted using FastStart High Fidelity PCR System (Roche), dNTP reagents mix (Bioline) with 60ng of DNA.

### PCR-based *FLG* enrichment with the Access Array 48.48 IFC

Fluidigm Access Array 48.48 IFC microfluidic chip enables 2304 PCR reactions to be performed simultaneously on a nanoscale level. Each chip can amplify a maximum of 48 DNA samples using 48 primer pair assays.

The *FLG* gene in each DNA sample was enriched and amplified via PCR using Access Array standard protocol. Briefly, 50 ng of DNA from each patient sample was pre-mixed with 1 μM of unique barcoding primer pair (Illumina), FastStart High Fidelity PCR reagents (Roche) and dNTP reagents mix (Bioline) prior to loading on to the Access Array 48.48 IFC (Fluidigm). 48 DNA mixtures and 48 *FLG* primer assays were then added to individual inlet wells of the Access Array IFC according to manufacturer’s instructions (Supplemental Figure 1). Thermocycling of primed and loaded IFC for PCR amplification and barcoding was conducted on the Biomark HD PCR machine (Fluidigm).

### Roche 454 GS FLX+ and Illumina MiSeq sequencing of *FLG* Fluidigm products

PCR product pool for each DNA sample was harvested from the IFC and individually quantified and checked for quality using Agilent DNA 1000 chip run on Bioanalyzer (Agilent). After determining correct size range of products, 1µL of PCR products from each sample (up to a maximum of 48 samples) were pooled and purified using Agencourt^®^ AMPure^®^ XP Reagent Beads (Beckman Coulter Genomics) according to the instructions provided in the Access Array Manual. Two or four pools of purified products were then further combined into a single pool containing 96 samples or 192 samples respectively before sequencing on Roche 454 GS FLX+ or Illumina MiSeq (v3 2 × 300bp or v2 2 × 250 bp read mode).

### Bioinformatics analysis of 454 and MiSeq

Raw sequence files were processed using a custom pipeline implemented in Pipeline Pilot (www.accelrys.com). Input reads were mapped to a *FLG* reference sequence containing 12 repeats that included the intragenic CNVs, introns, 5’ and 3’ UTRs that is not currently available as an NCBI RefSeq (Sandilands et al. 2007). We used BWA-MEM (version 0.7.10-r789) with the following parameters: gap opening penalty = 8, gap extension penalty = 2, mismatch penalty = 4, match score = 1 and minimum score threshold = 100. The mapped reads are stored as indexed BAM files which are then processed with GATK toolkit (version v3.4-46-gbc02625, Java version 1.7.0_75). *HaplotypeCaller* module was used to calculate SNVs and indels for each sample separately. Settings for *HaplotypeCaller* were as follows: minimum variant quality = 10, minimum pass quality = 50, maximum deletion length =12 and coverage was downsampled to 2500 in high-coverage regions. The individual gVCF files for each sample can then be collectively processed with the *GenotypeGVCFs* module to produce a joint VCF report for multiple samples (same quality cut-offs). This approach is flexible and scalable and increases the sensitivity of SNV detection (McKenna et al. 2010).

The pipeline returns a VCF report. The user can optionally reverse complement and convert the variants to GRCh37 or GRCh38 coordinates. Variants identified in the 8^2^ or 10^2^ repeat were not mapped to these reference coordinates. The VCF files can be annotated with the SnpEff tool (version 4.2) to annotate the SNVs and assess their impact. SnpEff was run with default options with the exception of lof.deleteProteinCodingBases which was set to 0.50.

### PCR and Sanger-sequencing of *FLG*

We comprehensively amplified the entire *FLG* open reading frame as previously described (Sandilands et al. 2007; Chen et al. 2011). PCR products were Sanger sequenced using the chain-terminator fluorescently-labelled dideoxyribonucleotidetriphosphate method (BigDye V3, ABI) and subjected to capillary electrophoresis on a ABI Prism^®^ 377 DNA Sequencer in order to determine the sequence of each PCR product (Smith et al., 2006; Chen et al., 2011). Electropherograms were analyzed using the Lasergene suite of sequencing analysis programs (DNASTAR). Sanger sequencing was also used to validate all *FLG* LoF variants identified by NGS using previous published primers (Sandilands et al. 2007; Chen et al. 2011).

Repeat 8 and repeat 10 regions in *FLG* exon 3 were amplified by PCR reactions (20 μL) using the Expand High Fidelity^PLUS^ PCR System (Roche) with 100 ng of genomic DNA, 1.5 U of enzyme, 1× Buffer 2, dNTPs (250 µM), 0.5 µM each of the forward and reverse primers and 4% v/v DMSO. PCR products were separated on a 0.9% w/v agarose gel and CNVs distinguished by their different sized products. The primers used for repeat 8 specific PCR were 5’-CCCAGGACAAGCAGGAACT-3’ and 5’- GCTTCATGGTGATGCGACCA-3’ (Sandilands et al. 2007). The primers used for repeat 10 specific PCR were 5’- GGGCCCAGGACAAGCAGGAAC-3’ (in-house designed) and 5’- CTGCACTACCATAGCTGCC-3’ (Sandilands et al. 2007).

### Statistical analysis of *FLG*

Allele frequencies were analyzed with Fisher’s exact test using a genotype based model with false discovery rates provided. Population allele frequency control data was extracted from ExAC Database (version 0.3.1) (Lek et al. 2016) using the East Asia subset exome data for comparison with Singaporean Chinese patients. South Asia and East Asia exome subset allele frequency data was again extracted from ExAC database and provided as control data for mutations identified in the Singapore Indian and Malay patients. However, because it was not clear if the South Asia and East Asia ethnicities in ExAC were a good matched control for the Indian and Malay ethnicities in this study, Fisher’s exact test was not performed to avoid any over interpretation of results. Analysis was performed in R (v3.3.2) with default settings for the Fisher’s exact test function.

## DATA ACCESS

All sequencing files were submitted to NCBI SRA and are publicly available under BioProject ID “PRJNA360024”. Accession codes for the individual samples are included in Supplemental Table 11.

## MATERIALS AND CORRESPONDENCE

Correspondence should be addressed to: J.E.A.C.

## ACKNOWLEDGEMENTS

We thank all the patients for donating to this study, the research coordinators at National Skin Centre, especially Nancy Liew and Veron Lu, for diligently collecting samples. Genome Institute of Singapore (GIS) Sequencing platform, A*STAR for their assistance. MiSeq assay development and quality control was checked with the kind assistance of Christopher Wong, Hui Mann Seah and Zhengzhong Qu, Polaris, GIS, A*STAR. This work was funded by A*STAR SPF grants for basic and translational skin research (IAF SPF 2013/004; IAF SPF 2013/005) to J.E.A.C., E.B.L., J.N.F., S.L.I.J.D. and J.L., and A*STAR SPF genetic orphan diseases (IAF SPF 2012/005) to X.F.C.C.W.

## CONTRIBUTIONS

J.E.A.C. and W.H.I.M designed the study components and planned the study. X.F.C.C.W., J.N.F., S.L.I.J.D. A.S.L.T., R.L.H. and H.J.C. designed and conducted all experiments under the guidance of E.B.L., J.L. and J.E.A.C. Patient samples collection was coordinated and supervised by M.B.Y.T. at National Skin Centre. X.F.C.C.W., S.L.I.J.D., E.B.L., and J.E.A.C. wrote the manuscript with inputs from all authors.

## REFERENCES

Artuso R, Fallerini C, Dosa L, Scionti F, Clementi M, Garosi G, Massella L, Epistolato MC, Mancini R, Mari F et al. 2012. Advances in Alport syndrome diagnosis using next-generation sequencing. European journal of human genetics : EJHG 20(1): 50–57.

Balzer S, Malde K, Jonassen I. 2011. Systematic exploration of error sources in pyrosequencing flowgram data. Bioinformatics 27(13): i304–309.

Berg JS, Evans JP, Leigh MW, Omran H, Bizon C, Mane K, Knowles MR, Weck KE, Zariwala MA. 2011. Next generation massively parallel sequencing of targeted exomes to identify genetic mutations in primary ciliary dyskinesia: implications for application to clinical testing. Genetics in medicine : official journal of the American College of Medical Genetics 13(3): 218–229.

Brown SJ, Kroboth K, Sandilands A, Campbell LE, Pohler E, Kezic S, Cordell HJ, McLean WHI, Irvine AD. 2012. Intragenic copy number variation within filaggrin contributes to the risk of atopic dermatitis with a dose-dependent effect. Journal of Investigative Dermatology 132(1): 98–104.

Brown SJ, McLean WHI. 2012. One remarkable molecule: filaggrin. Journal of Investigative Dermatology 132(3 Pt 2): 751–762.

Brown SJ, Relton CL, Liao H, Zhao Y, Sandilands A, Wilson IJ, Burn J, Reynolds NJ, McLean WHI, Cordell HJ. 2008a. Filaggrin null mutations and childhood atopic eczema: a population-based case-control study. The Journal of allergy and clinical immunology 121(4): 940–946.e943.

Brown SJ, Sandilands A, Zhao Y, Liao H, Relton CL, Meggitt SJ, Trembath RC, Barker JNWN, Reynolds NJ, Cordell HJ et al. 2008b. Prevalent and low-frequency null mutations in the filaggrin gene are associated with early-onset and persistent atopic eczema. The Journal of investigative dermatology 128(6): 1591–1594.

Chen H, Common JEA, Haines RL, Balakrishnan A, Brown SJ, Goh CSM, Cordell HJ, Sandilands A, Campbell LE, Kroboth K et al. 2011. Wide spectrum of filaggrin-null mutations in atopic dermatitis highlights differences between Singaporean Chinese and European populations. The British journal of dermatology.

Cingolani P, Platts A, Wang le L, Coon M, Nguyen T, Wang L, Land SJ, Lu X, Ruden DM. 2012. A program for annotating and predicting the effects of single nucleotide polymorphisms, SnpEff: SNPs in the genome of Drosophila melanogaster strain w1118; iso-2; iso-3. Fly 6(2): 80–92.

Clark AG, Hubisz MJ, Bustamante CD, Williamson SH, Nielsen R. 2005. Ascertainment bias in studies of human genome-wide polymorphism. Genome research 15(11): 1496–1502.

Eaaswarkhanth M, Xu D, Flanagan C, Rzhetskaya M, Hayes MG, Blekhman R, Jablonski N, Gokcumen O. 2016. Atopic Dermatitis Susceptibility Variants In Filaggrin Hitchhike Hornerin Selective Sweep. Genome biology and evolution.

Fallon PG, Sasaki T, Sandilands A, Campbell LE, Saunders SP, Mangan NE, Callanan JJ, Kawasaki H, Shiohama A, Kubo A et al. 2009. A homozygous frameshift mutation in the mouse Flg gene facilitates enhanced percutaneous allergen priming. Nat Genet 41(5): 602–608.

Gan SQ, McBride OW, Idler WW, Markova N, Steinert PM. 1990. Organization, structure, and polymorphisms of the human profilaggrin gene. Biochemistry 29(40): 9432–9440.

Goudie DR, DʼAlessandro M, Merriman B, Lee H, Szeverényi I, Avery S, Oʼconnor BD, Nelson SF, Coats SE, Stewart A et al. 2011. Multiple self-healing squamous epithelioma is caused by a disease-specific spectrum of mutations in TGFBR1. Nature genetics.

Henry J, Toulza E, Hsu CY, Pellerin L, Balica S, Mazereeuw-Hautier J, Paul C, Serre G, Jonca N, Simon M. 2012. Update on the epidermal differentiation complex. Frontiers in bioscience 17: 1517–1532.

Irvine AD, McLean WHI. 2006. Breaking the (un)sound barrier: filaggrin is a major gene for atopic dermatitis. The Journal of investigative dermatology 126(6): 1200–1202.

Irvine AD, McLean WHI, Leung DYM. 2011. Filaggrin mutations associated with skin and allergic diseases. The New England journal of medicine 365(14): 1315–1327.

Kono M, Nomura T, Ohguchi Y, Mizuno O, Suzuki S, Tsujiuchi H, Hamajima N, McLean WHI, Shimizu H, Akiyama M. 2014. Comprehensive screening for a complete set of Japanese-population-specific filaggrin gene mutations-Kono-2014-Allergy-Wiley Online Library. Allergy 69(4): 537–540.

Lange V, Bohme I, Hofmann J, Lang K, Sauter J, Schone B, Paul P, Albrecht V, Andreas JM, Baier DM et al. 2014. Cost-efficient high-throughput HLA typing by MiSeq amplicon sequencing. BMC genomics 15: 63.

Lek M, Karczewski KJ, Minikel EV, Samocha KE, Banks E, Fennell T, O'Donnell-Luria AH, Ware JS, Hill AJ, Cummings BB et al. 2016. Analysis of protein-coding genetic variation in 60,706 humans. Nature 536(7616): 285–291.

Li H. 2013. Aligning sequence reads, clone sequences and assembly contigs with BWA-MEM. In arXivorg. Oxford University Press, arXiv:1303.3997v2 [q-bio.GN].

Margolis DJ, Apter AJ, Mitra N, Gupta J, Hoffstad O, Papadopoulos M, Rebbeck TR, MacCallum S, Campbell LE, Sandilands A et al. 2013. Reliability and validity of genotyping filaggrin null mutations. Journal of dermatological science 70(1): 67–68.

Margolis DJ, Gupta J, Apter AJ, Hoffstad O, Papadopoulos M, Rebbeck TR, Wubbenhorst B, Mitra N. 2014. Exome Sequencing of Filaggrin and Related genes in African-American Children with Atopic Dermatitis. Journal of Investigative Dermatology.

McAleer MA, Irvine AD. 2013. The multifunctional role of filaggrin in allergic skin disease. J Allergy Clin Immunol 131(2): 280–291.

McKenna A, Hanna M, Banks E, Sivachenko A, Cibulskis K, Kernytsky A, Garimella K, Altshuler D, Gabriel S, Daly M et al. 2010. The Genome Analysis Toolkit: a MapReduce framework for analyzing next-generation DNA sequencing data. Genome research 20(9): 1297–1303.

Nagao K, Segre JA. 2015. “Bringing Up Baby” to Tolerate Germs. Immunity 43(5): 842–844.

Ng SB, Turner EH, Robertson PD, Flygare SD, Bigham AW, Lee C, Shaffer T, Wong M, Bhattacharjee A, Eichler EE et al. 2009. Targeted capture and massively parallel sequencing of 12 human exomes. Nature 461(7261): 272–276.

Odhiambo JA, Williams HC, Clayton TO, Robertson CF, Asher MI, Group IPTS. 2009. Global variations in prevalence of eczema symptoms in children from ISAAC Phase Three. The Journal of allergy and clinical immunology 124(6): 1251–1258.e1223.

Oranje AP, Glazenburg EJ, Wolkerstorfer A, de Waard-van der Spek FB. 2007. Practical issues on interpretation of scoring atopic dermatitis: the SCORAD index, objective SCORAD and the three-item severity score. Br J Dermatol 157(4): 645–648.

Palmer CNA, Irvine AD, Terron-Kwiatkowski A, Zhao Y, Liao H, Lee SP, Goudie DR, Sandilands A, Campbell LE, Smith FJD et al. 2006. Common loss-of-function variants of the epidermal barrier protein filaggrin are a major predisposing factor for atopic dermatitis. Nature genetics 38(4): 441–446.

Robinson JT, Thorvaldsdottir H, Winckler W, Guttman M, Lander ES, Getz G, Mesirov JP. 2011. Integrative genomics viewer. Nature biotechnology 29(1): 24–26.

Ruderfer DM, Hamamsy T, Lek M, Karczewski KJ, Kavanagh D, Samocha KE, Exome Aggregation C, Daly MJ, MacArthur DG, Fromer M et al. 2016. Patterns of genic intolerance of rare copy number variation in 59,898 human exomes. Nat Genet 48(10): 1107–1111.

Sandilands A, O’Regan GM, Liao H, Zhao Y, Terron-Kwiatkowski A, Watson RM, Cassidy AJ, Goudie DR, Smith FJ, McLean WH et al. 2006. Prevalent and rare mutations inthe gene encoding filaggrin cause ichthyosis vulgaris and predispose individuals to atopic dermatitis. J Invest Dermatol 126(8): 1770–1775.

Sandilands A, Sutherland C, Irvine AD, McLean WHI. 2009. Filaggrin in the frontline: role in skin barrier function and disease. Journal of cell science 122(Pt 9): 1285–1294.

Sandilands A, Terron-Kwiatkowski A, Hull PR, OʼRegan GM, Clayton TH, Watson RM, Carrick T, Evans AT, Liao H, Zhao Y et al. 2007. Comprehensive analysis of the gene encoding filaggrin uncovers prevalent and rare mutations in ichthyosis vulgaris and atopic eczema. Nature genetics 39(5): 650–654.

Sasaki T, Kudoh J, Ebihara T, Shiohama A, Asakawa S, Shimizu A, Takayanagi A, Dekio I, Sadahira C, Amagai M et al. 2008. Sequence analysis of filaggrin gene by novel shotgun method in Japanese atopic dermatitis. J Dermatol Sci 51(2): 113–120.

Schwarz JM, Cooper DN, Schuelke M, Seelow D. 2014. MutationTaster2: mutation prediction for the deep-sequencing age. Nature methods 11(4): 361–362.

Scott CA, Plagnol V, Nitoiu D, Bland PJ, Blaydon DC, Chronnell CM, Poon DS, Bourn D, Gárdos L, Császár A et al. 2012. Targeted Sequence Capture and High-Throughput Sequencing in the Molecular Diagnosis of Ichthyosis and Other Skin Diseases. Journal of Investigative Dermatology 133(2): 573–576.

Smith FJD, Irvine AD, Terron-Kwiatkowski A, Sandilands A, Campbell LE, Zhao Y, Liao H, Evans AT, Goudie DR, Lewis-Jones S et al. 2006. Loss-of-function mutations in the gene encoding filaggrin cause ichthyosis vulgaris. Nature genetics 38(3): 337–342.

Strasser B, Mlitz V, Hermann M, Rice RH, Eigenheer RA, Alibardi L, Tschachler E, Eckhart L. 2014. Evolutionary origin and diversification of epidermal barrier proteins in amniotes. Molecular biology and evolution 31(12): 3194–3205.

Tay Y-K, Kong K-H, Khoo L, Goh C-L, Giam Y-C. 2002. The prevalence and descriptive epidemiology of atopic dermatitis in Singapore school children. The British journal of dermatology 146(1): 101–106.

Thyssen JP, Bikle DD, Elias PM. 2014. Evidence That Loss-of-Function Filaggrin Gene Mutations Evolved in Northern Europeans to Favor Intracutaneous Vitamin D3 Production. Evolutionary biology 41(3): 388–396.

Thyssen JP, Thuesen B, Huth C, Standl M, Carson CG, Heinrich J, Krämer U, Kratzsch J, Berg ND, Menné T et al. 2012. Skin barrier abnormality caused by filaggrin (FLG) mutations is associated with increased serum 25-hydroxyvitamin D concentrations. Journal of Allergy and Clinical Immunology 130(5): 1204–1207.e1202.

van denOord RAHM, Sheikh A. 2009. Filaggrin gene defects and risk of developing allergic sensitisation and allergic disorders: systematic review and meta-analysis. BMJ (Clinical research ed) 339: b2433.

Wells RS, Kerr CB. 1966. Clinical features of autosomal dominant and sex-linked ichthyosis in an English population. British medical journal 1(5493): 947–950.

